# Genomic Selection in Rubber Tree Breeding: A Comparison of Models and Methods for dealing with G × E

**DOI:** 10.1101/603662

**Authors:** L.M. Souza, F.R. Francisco, P.S. Gonçalves, E.J. Scaloppi Junior, V. Le Guen, R. Fritsche-Neto, A.P Souza

**Affiliations:** Molecular Biology and Genetic Engineering Center (CBMEG), University of Campinas (UNICAMP), Campinas, SP, Brazil; Center of Rubber Tree and Agroforestry Systems, Agronomic Institute (IAC), Votuporanga, SP, Brazil; Centre de Coopération Internationale en Recherche Agronomique pour le Développement (CIRAD) UMR AGAP, Montpellier, Hérault, France; Departamento de Genética, Escola Superior de Agricultura “Luiz de Queiroz” Universidade de São Paulo (ESALQ/USP), Piracicaba, SP, Brazil; Department of Plant Biology, Biology Institute, University of Campinas (UNICAMP) UNICAMP, Campinas, SP, Brazil

**Keywords:** *Hevea brasiliensis*, Breeding, Multienvironment, Single-nucleotide, Genotyping

## Abstract

Several genomic prediction models incorporating genotype × environment (G×E) interactions have recently been developed and used in genomic selection (GS) in plant breeding programs. G×E interactions decrease selection accuracy and limit genetic gains in plant breeding. Two genomic data sets were used to compare the prediction ability of multi-environment G×E genomic models and two kernel methods (a linear kernel (genomic best linear unbiased predictor, GBLUP) (GB) and a nonlinear kernel (Gaussian kernel, GK)) and prediction accuracy (PA) of five genomic prediction models: (1) one without environmental data (BSG); (2) a single-environment, main genotypic effect model (SM); (3) a multi-environment, main genotypic effect model (MM); (4) a multi-environment, single variance GxE deviation model (MDs); and (5) a multi-environment, environment-specific variance GxE deviation model (MDe). We evaluated the utility of GS with 435 rubber tree individuals in two sites and genotyped the individuals with genotyping-by-sequencing (GBS) of single-nucleotide polymorphisms (SNPs). Prediction models were estimated for diameter (DAP) and height (AP) at different ages, with a heritability ranging from 0.59 to 0.75 for both traits. Applying the model (BSG, SM, MM, MDs, and MDe) and kernel method (GBLUP and GK) combinations to rubber tree data showed that models with the nonlinear GK and linear GBLUP kernel had similar PAs. Multi-environment models were superior to single-environment genomic models regardless the kernel (GBLUP or GK), suggesting that introducing interactions between markers and environmental conditions increases the proportion of variance explained by the model and, more importantly, the PA. In the best scenario (well-watered (WW / GK), an increase of 6.7 and 8.7 fold of genetic gain can be obtained for AP and DAP, respectively, with multi-environment GS (MM, MDe and MDS) than by conventional genetic breeding model (CBM). Furthermore, GS resulted in a more balanced selection response in DAP and AP and if used in conjunction with traditional genetic breeding programs will contribute to a reduction in selection time. With the rapid advances in and declining costs of genotyping methods, balanced against the overall costs of managing large progeny trials and potential increased gains per unit time, we are hopeful that GS can be implemented in rubber tree breeding programs.

## 1 Introduction

Generally, the rubber tree breeding program is characterized by breeding cycles of 25-30 years and includes the production of crosses, evaluation, and selection of field progeny, and propagation of selected superior material (Gonçalves et al., 2006). Compared to animal and annual crop breeding, forest tree breeding is still in its infancy, and the most advanced programs are in their third or fourth cycle of breeding, with very little differentiation of the bred populations from natural populations (Isik, 2014). Rubber tree breeding programs are complex and costly because the large size of trees requires experiments over large tracts of land to test progeny, and the progeny tests are expensive to establish, manage over many years, and evaluate via measurement.

The main objective of rubber tree breeding is the development of early selection methods that support the accurate prediction of mature phenotypes at a younger stage and are therefore important for shortening breeding cycles and, in the end, improving the cost efficiency of such breeding programs. *Hevea* breeding needs to significantly reduce the time taken to derive a clone, Priyadarshan (2017) proposed two strategies: (1) to cut short the breeding steps being followed by conventional means and (2) to inculcate genomics into breeding programmes specially to identify high-yielding genotypes in half-sibs, full-sibs and poly-cross seedlings during juvenile stage that can minimize both space and time.

Traditional plant breeding programs depend mainly on phenotypes being evaluated in various environments; selection and recombination are based solely on the resulting data plus pedigree information, when available. Genomic selection (GS), a new approach using whole-genome molecular markers, has the potential to quickly improve complex traits with low heritability, significantly reduce the cost of the line and hybrid development and increase grain production in less time to improve quantitative traits in large plant breeding populations (Meuwissen et al. 2001).

Genomic prediction combines marker data with phenotypic and pedigree data in an attempt to increase the accuracy of the prediction of breeding and genotypic values. The method depends on dense genome-wide marker coverage to produce genomic estimated breeding values (GEBVs) from an ensemble analysis of all markers.

According to Lorenz et al. (2011), the accuracy of GS, which is measured as the correlation between the GEBVs and true breeding values, is affected by the relationship between the training and test sets, the number of individuals in the training set, linkage disequilibrium (LD) between markers and quantitative trait loci (QTLs), the distribution of underlying QTL effects, the statistical method used to estimate the GEBVs, and the trait heritability.

GS was proposed by Meuwissen et al. (2001) and has received increasing interest from forest tree breeders. The initial experimental reports in Pinus and Eucalyptus (Resende et al., 2012a,b) demonstrated the encouraging prospects of this new method and have since confirmed the potential for GS in conifers, pines and eucalypts (Zapata-Valenzuela et al., 2013; Lima, 2014; El-Dien et al., 2015; Ratcliffe et al., 2015; Bartholome et al., 2016; Isik et al., 2016), further supporting the potential for GS to accelerate the breeding of forest trees.

In the rubber tree breeding program, pedigree-based analysis has been widely used to evaluate field experiments, estimate genetic parameters, and predict breeding values (Furlani et al., 2005). However, due to the decreasing costs of genotyping thousands or millions of markers and the increasing costs of phenotyping (Krchov and Bernardo, 2015), GS is arising as an alternative genome-wide marker-based method to predict future genetic responses.

Appropriate GS methods provide accurate predictions even for untested genotypes, allowing considerable progress in breeding programs by reducing the number of field-tested genotypes and, consequently, the costs of phenotyping (Krchov and Bernardo, 2015). The benefits of GS are more evident when traits are difficult, time-consuming, expensive to measure, and several environments need to be evaluated.

The objective of this paper was to evaluate the predictive capacity of GS implementation in rubber trees using linear and nonlinear kernel methods and the performance of such prediction when including GxE interactions in each of the four models described by Bandeira et al. (2017). Thus, for all data sets, we fitted models with a linear kernel using the genomic best linear unbiased predictor (GBLUP) (GB) or nonlinear Gaussian kernel (GK) with a bandwidth parameter estimated according to (Pérez-Elizalde et al., 2015). We also compared the prediction accuracy (PA) of the two kernel regression methods for the four models. The models included a single-environment, main genotypic effect model (SM), a multi-environment, main genotypic effect model (MM) (Jarquin et al., 2014), a multi-environment, single variance G×E deviation model (MDs) (Jarquin et al., 2014) and a multiple-environment, environment-specific variance G×E deviation model (MDe) (Lopez-Cruz et al., 2015).

To the best of our knowledge, this is the first attempt to apply the GS technique in a rubber tree breeding program. The development of robust methods enables the implementation of GS in routine evaluations to accelerate genetic progress.

## 2 Materials and methods

### 2.1 Population and phenotypes

The data set included 435 samples, consisting of 252 F1 hybrids derived from a cross between PR255 × PB217 from the Michelin Ltda. Breeding program (Souza et al., 2013), 146 F1 hybrids derived from open pollination between the genotypes GT1 and RRIM701 (Conson et al., 2018), 37 genotypes of GT1 × PB235 crosses and four testers (GT1, PB235, RRIM701 and the commercial clone RRIM600) from the Agronomic Institute Campinas breeding program. The parents of two-parent families are not related to each other, except PB217 and PB235 who are half-siblings.

Two traits were analyzed: (i) height of the trees (AP), taken at the insertion of the highest leaf into the trunk, and (ii) circumference of the trunk (DAP), measured 1 m above the soil (data available in Souza et al., 2013; Conson et al., 2018), in two periods: low water (LW) and well-watered (WW) (Supplementary Table 1).

### 2.2 Genotypic data and single nucleotide polymorphism (SNP) calling

Genomic DNA was extracted according to Souza et al. (2013) and Conson et al. (2018). Genotyping-by-sequencing (GBS) library preparation and sequencing were performed as described by Elshire et al. (2011). Genome complexity was reduced by digesting individual genomic DNA samples with *EcoT22I*, a methylation-sensitive restriction enzyme, and 96 samples per sequencing lane. The resulting fragments from each sample were directly ligated to a pair of enzyme-specific adapters and combined into pools. PCR amplification was carried out to generate the GBS libraries, which were sequenced with the Illumina platform (Illumina Inc., USA).

The raw data were processed, and SNP calling was performed using TASSEL 5.0 (Glaubitz et al., 2014). Initially, the FASTQ files were demultiplexed according to the assigned barcode. The reads from each sample were trimmed, and the tags were identified using the following parameters: a kmer length of 64 bp, minimum quality score within the barcode and read length of 20, minimum kmer length of 20 and a minimum count of reads for a tag of 6. All sequence tags from each sample were aligned to the reference rubber tree genome (Tang et al., 2016) with Bowtie 2 (Langmead and Salzberg, 2012) using the very sensible option. SNP calling was performed using the TASSEL 5 GBSv2 pipeline (Glaubitz et al., 2014) and filtered with snpReady software (Granato and Fritsche-Neto, 2018). The following criteria were used: missing data of 20% and minor allele frequency (MAF) greater than or equal to 5% (MAF of 0.05). Only biallelic SNPs were maintained using the software VCFtools (Danecek et al., 2011). After filtering, missing data were imputed using snpReady software (Granato and Fritsche-Neto, 2018).

### 2.3 GS analysis

For each character, the phenotypic analysis was carried out jointly for all years of evaluation using the mixed model approach.

Prediction based on genomic relationships and predictive ability assessment was performed using a relationship matrix-based approach for genomic prediction (Habier et al., 2007); the matrix G was the central object denoting the genomic relationship matrix. Two kernel methods were used: the linear kernel (GBLUP, GB) method used by Jarquin et al. (2014) and Lopez-Cruz et al. (2015) and the nonlinear kernel (GK) method proposed by Cuevas et al. (2016). The matrix for the GB and GK methods was obtained with the function *G.matrix* in snpReady software (Granato and Fritsche-Neto, 2018). Statistical models for genomic predictions taking genotype × environment (G×E) interactions into account (Jarquin et al., 2014; Lopez-Cruz et al., 2015) combine genetic information from molecular markers or from pedigrees (Pérez-Rodríguez et al., 2015) with environmental covariates, while the López-Cruz model decomposes the marker effect across all environments and the interaction for each specific environment.

The GS models implemented with arrays of GK and GB pedigrees for the AP and DAP traits were implemented in *breedR* software (Munõz and Sanchez, 2017); using frequentist statistics with the function *remlf90, em* method, 5 folds and 5 repetitions, the training population (TRN) was created with 4 folds, whereas the test population (TST) was created with one fold. The PA was obtained from the correlation between the predicted *BLUPs* and the observed *BLUPs.*

For AP and DAP, five statistical prediction models were fitted to all data sets to study their PA using random cross-validation (CV) schemes. The main objective was to compare the prediction ability of the two proposed multi-environment G×E genomic models.

The PA of the two kernel regression methods was also compared for single environments and multi-environments: a single-environment, main genotypic effect model (SM), a multi-environment, main genotypic effect model (MM) (Jarquin et al., 2014), a multi-environment, single variance G×E deviation model (MDs) (Jarquin et al., 2014) and a multi-environment, environment-specific variance G×E deviation model (MDe) (Lopez-Cruz et al., 2015). The SM, MM, MDs, and MDe models fitted with the GB and GK methods were used on the entire data sets for all the traits, and the phenotypic data were centered and standardized. These analyses were performed to derive estimates of variance components. The following variance components resulting from the residual effects, main genetic effect, and genetic environment-specific effects of the four models described above for the trait (LW – low water, and WW – well-watered) data sets were computed. All models were fitted with GxE interactions using the software BGGE (Granato et al., 2018).

### 2.4 Assessing PA by random cross-validation (CV) GxE

The PA of the SM model-method combinations was evaluated with 80% of the hybrids comprising the TRN set, the remaining 20% of the individuals comprising the TST set and none of the lines to be predicted in the TST set in the TRN set using 5 random partitions arranged in 5 folds with 100 random partitions each. This procedure was performed separately in each environment, namely, LW and WW, and the SM models were fitted separately for each environment.

In the multi-environment models, the PA of the model-method combinations was generated using two different CV designs (Burgueño et al., 2012). The random CV 1 design (CV1) assumes that newly developed lines have not been evaluated in any environment; in this case, 20% of the lines were not observed (not phenotyped) in all the environments and had to be predicted. The random CV 2 design (CV2) simulates lines that are tested in incomplete field trials, where some lines are evaluated in some environments but not included in other environments.

All the parameters of the models, including the variance components resulting from residual effects, main genetic effects, G × E interaction effects, and environment-specific effects, were re-estimated from the TRN data in each of the 50 random TRN-TST partitions, and models were fitted to the TRN data set. PA was assessed by computing Pearson’s product-moment correlation between predictions and phenotypes in the TST data set within environments.

### 2.5 Expected genetic gain (EGG)

Selection gain was estimated in two ways: the classic way for rubber tree breeding using the breeder’s equation and phenotypic data and with information from the SNPs obtained via GS.

### 2.6 EGGc – selection gain using only phenotypic information

The genetic gains obtained by a classical breeding cycle were estimated under the assumption that the time of selection is ten years (EGGc), representing the minimum time required to make the crosses, obtain seeds, and evaluate and select the progenies at a small scale:

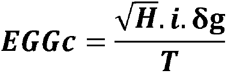

EGGc= Expected Gain of Classic Breeding Selection

H= Broad heritability

i= Selected individuals

*δ*g= Additive genetic standard deviation

### 2.7 EGGgs – Selection gain using molecular marker information

The simulation of breeding cycles using GS was based on the EGGgs equation assuming a time of 3 years for each selection cycle, representing the time required for crossing, seed selection and selection of the best individuals via molecular markers.

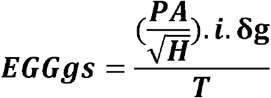

EGGgs = Expected Gain of Selection with Genomic Selection

PA= Prediction accuracy

H= Broad heritability

i= Selected individuals

*δ*g= Additive genetic standard deviation

## 3 Results

### 3.1 SNP calling

We started with 435 genotypes, but three genotypes were replicates and thus merged, and the function sweep removed 27 individuals based on criteria of <0.05 (for quality purposes), leaving 411 genotypes. After analysis, a total of 259.224 million reads of sequence data were obtained, of which 69.8% were good barcoded reads. The overall alignment rate of these reads to the rubber tree reference genome (Tang et al., 2016) was 83.70%, and 23.1% were aligned exactly one time.

A total of 107.294 SNPs were identified. After excluding markers (1) with more than 20% missing data, (2) with a MAF ≤ 0.05 or (3) SNPs with more than two alleles, the whole dataset was reduced to 30.546 SNPs.

### 3.2 Estimates of genetic parameters using SNP genotyping

With the genotyped SNPs, the population structure was assessed using a principal component analysis (PCA), and the plots indicated that the 411 genotypes fell into two major clusters (Supplementary Figure 1), which mainly contained hybrids derived from PR255 × PB217 and hybrids derived from GT1 x RRIM701 and of GT1 × PB235 crosses. The first two PCs explained 19.51% and 2.18% of the total variance, respectively, clearly splitting the groups along the x and y-axes.

### 3.3 Descriptive statistics

The genetic correlations between DAP and AP in both environments, namely, LW and WW, were positive and significant, ranging from 0.72 (AP-LW × DAP-WW) to 0.99 (DAP × DAP-LW)

To assess how much the phenotypic variation is genetically controlled and thus efficient for GS, we first estimated the broad-sense heritability (H) of DAP and AP. The heritability ranged from 0.60 to 0.75 for both traits, and when we analyzed the variation separately in each environment, namely, LW and WW, the heritability varied between 0.33 and 0.34 for DAP and 0.41 and 0.42 for AP (Table 1).

**Table 1.**
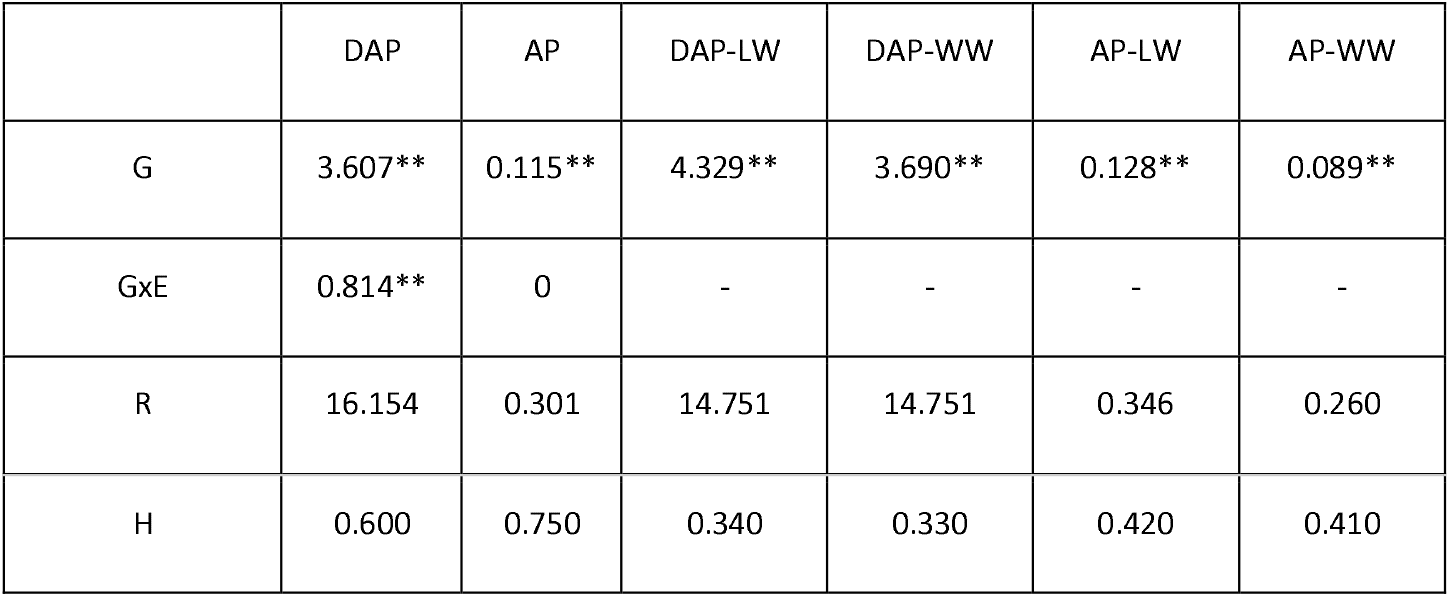
Phenotypic variation: heritability (H) of height (AP) and diameter (DAP), genotype × environment interaction (GxE), residual (R) and genetic main effect (G) in the low-water (LW) and well-watered (WW) environments considered together and alone, with p<.01 indicated by **

|  | DAP | AP | DAP-LW | DAP-WW | AP-LW | AP-WW |
| --- | --- | --- | --- | --- | --- | --- |
| G | 3.607** | 0.115** | 4.329** | 3.690** | 0.128** | 0.089** |
| GxE | 0.814** | 0 | - | - | - | - |
| R | 16.154 | 0.301 | 14.751 | 14.751 | 0.346 | 0.260 |
| H | 0.600 | 0.750 | 0.340 | 0.330 | 0.420 | 0.410 |

For all the evaluated characters, environmental variation, progeny and the interaction between these two factors were highly significant, indicating that the environments used were contrasting, there was genetic variability among genotypes, and these genotypes presented differential performance according to the environment, respectively. The coefficients of experimental variation obtained in the joint analysis were following those reported in the literature for the same characters, indicating that the experiments were performed with good precision. All components of the variance were nonzero, and estimates were obtained with reasonable accuracy, which was verified by the confidence intervals of these estimates (Table 1).

### 3.4 Estimates of variance components

Estimates of variance components for each of the GS models derived from the full data analysis are presented in Table 2.

**Table 2.** Estimates of different variance components for genomic selection (GS) models with no environmental effect; only the GB and GK (SG) matrices; the single-environment, main genotypic effect model (SM); the multi-environment, main genotypic effect model (MM); the multi-environment, single variance G×E deviation model (MDs); and the multi-environment, environment-specific variance G×E deviation model with the genomic best linear unbiased predictor (GBLUP) and Gaussian kernel (GK) for two traits: plant height (AP) and plant diameter (DAP).

|  | Component | DAP.GK | DAP.GB | AP.GK | AP.GB |
| --- | --- | --- | --- | --- | --- |
| BSG | G | 0.750 | 0.230 | 0.070 | 0.040 |
|  | E | 2.000 | 2.170 | 0.040 | 0.060 |
| SM | G (WW) | 1.102 | 1.101 | 0.062 | 0.069 |
|  | G (LW) | 2.528 | 2.691 | 0.105 | 0.140 |
|  | E (WW) | 1.532 | 1.773 | 0.037 | 0.052 |
|  | E (LW) | 2.628 | 3.164 | 0.047 | 0.065 |
| MM | G | 6.47 | 17.188 | 0.166 | 0.422 |
|  | E | 0.256 | 0.259 | 0.008 | 0.008 |
| MDs | G | 6.515 | 17.568 | 0.167 | 0.433 |
|  | E | 0.127 | 0.15 | 0.004 | 0.004 |
| MDe | G | 6.115 | 16.922 | 0.161 | 0.427 |
|  | E | 0.111 | 0.119 | 0.003 | 0.003 |

For the SM model of the two traits in each environment, the estimated residual variance components for the GK method were smaller than those for the GB method (Table 2), and in compensation, the variance in genetic effects in each environment was slightly more substantial for the SM-GB model-method combination than for the SM-GK combination. Both the genetic variance and phenotypic variance were more significant in LW than in WW.

The inclusion of the interaction term (G×E) when using the MM, MDe and MDs induced a more significant reduction in the estimated residual variance for both traits (DAP and AP) (Table 2). In the six model-method combinations, the residuals were even lower for DAP with the GK method, whereas for AP, the values were similar.

For the variance component associated with the genetic interaction effect, the values were, on average, 65% lower for the GK method than for the GB method.

### 3.5 Assessment of PA

The PA without environmental data (BSG) is shown in Figure 1. The estimated correlations between correlated phenotypes and predictions obtained in the CV test are shown in Figure 2 for the single-environment model (SM) and Figure 3 for the multi-environment models (MM, MDs and MDe).

**Figure 1.**
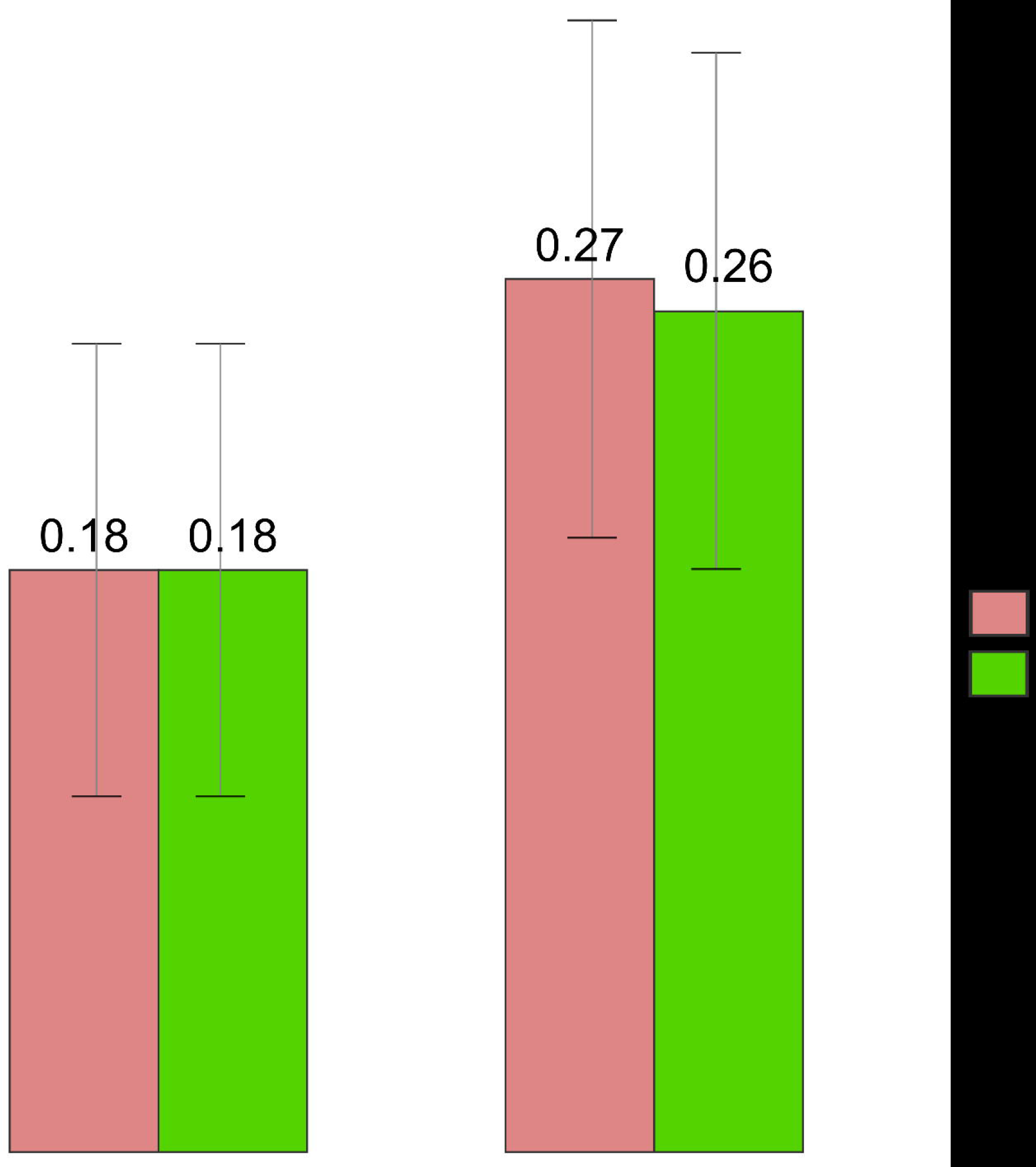
Correlation between phenotypes and predicted values (random CV partitions) with standard deviations for the GBLUP kernel model (GB) and GK model (GK) for the height of the trees (AP) and circumference of the trunk (DAP).

**Figure 2.**
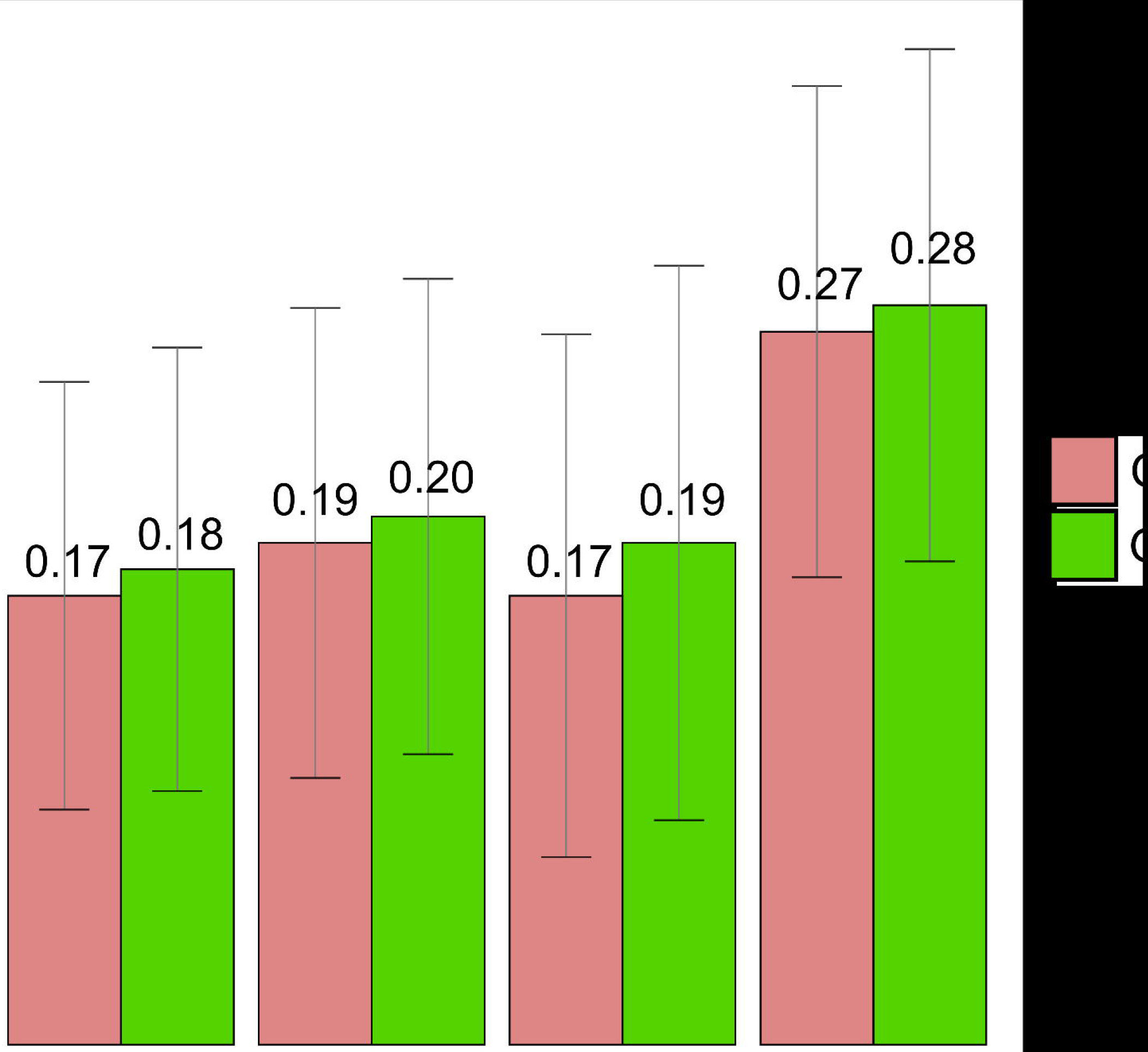
Correlation between phenotypes and predicted values (random CV partitions) with standard deviations for the single-environment, main genotypic effect model with the GBLUP kernel method (SM-GB) and the single-environment, main genotypic effect model with the GK method (SM-GK) for the height of the trees (AP) and circumference of the trunk (DAP) in low-water (LW) and well-watered (WW) environments.

**Figure 3.**
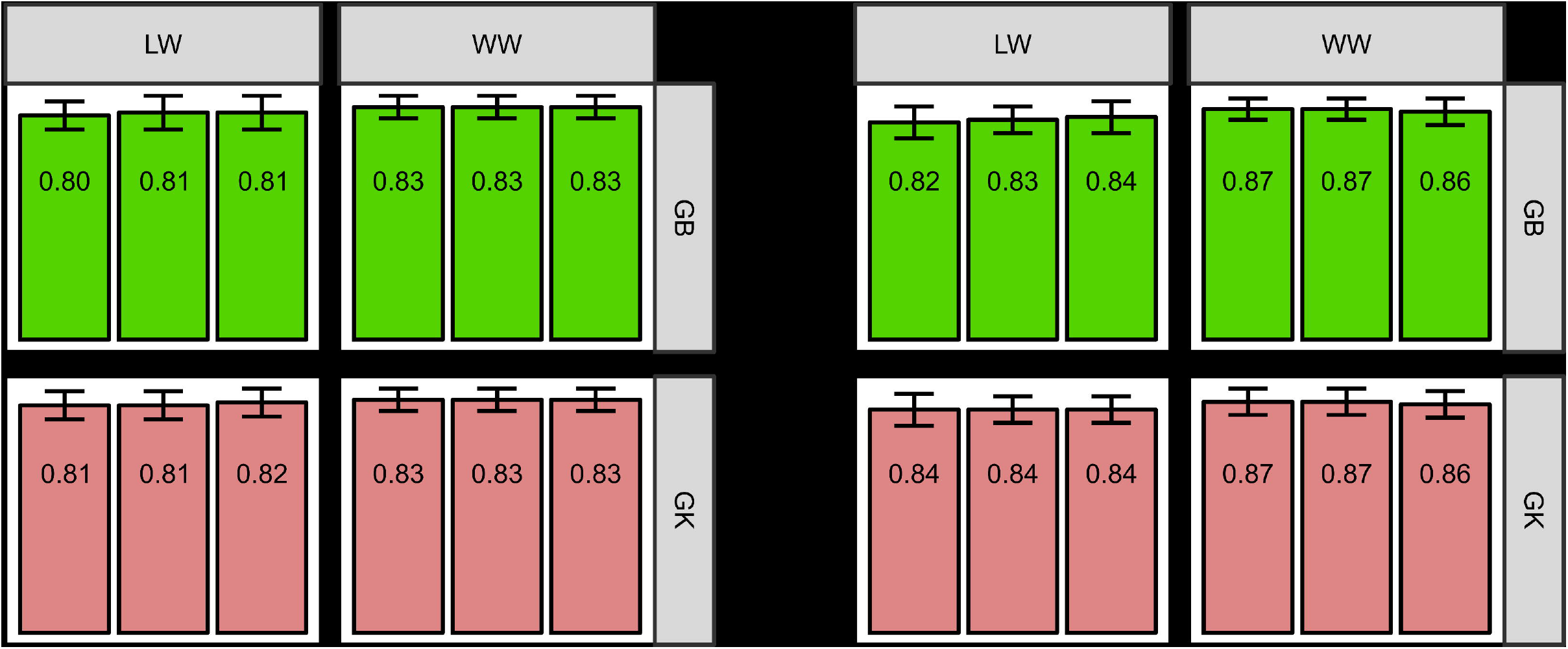
Prediction values for cross-validation (CV) partitions (CV2) and standard deviations for the multi-environment, genotypic effect model with the GBLUP kernel (MM-GB), multienvironment, main genotypic model with the Gaussian kernel (MM-GK), multi-environment, single variance G×E model with the GBLUP kernel (MDs-GB), multi-environment, single variance G×E model with the Gaussian kernel (MDs-GK), multi-environment, environment-specific variance G×E model with the GBLUP kernel (MDe-GB), and multi-environment, environment-specific variance G×E model with the Gaussian kernel (MDe-GK).

#### 3.5.1 PA without environmental data (BSG)

For DAP, the PA was not significantly different between the GB and GK models without multi-environmental data. The same result was obtained for GB and GK (0.18), whereas for DAP, there was a small difference between GK (0.26) and GB (0.27) (Figure 1).

#### 3.5.2 Single environment (SM)

For all traits, the random CV (CV1) applied to only one environment (LW or WW). The results for AP and DAP showed that the PA of the SM-GK model-method combination was higher than that for the SM-GB combination in both LW and WW. The PAs for AP in the LW conditions were 0.17 for SM-GB and 0.18 for SM-GK and in the WW environment were 0.19 for SM-GB and 0.20 for SM-GK. The results for DAP were 0.17 in LW for the SM-GB, and 0.19 for the SM-GK and in WW environment were 0.27 for SM-GB and 0.28 for SM-GK.

#### 3.5.3 Multi-environment (MM, MDe, and MDs)

The PA varied considerably between the CV1 and CV2 conditions, with an average PA 72.6% higher for CV2 than for CV1 (Supplementary Figure 3). Considering only random CV2, the PA was slightly higher in WW conditions for both characters (AP and DAP), ranging from 0.87 (MDs-GK-WW) to 0.80 (GK-MDe-LW).

The results obtained with the model-method combinations were very similar; generally, for the LW environment, the best model was GK, without any difference between the methods. In DAP-LW, the PA was 0.84, and for AP-LW, it ranged from 0.81 (MM) to 0.83 (MDe and MDs) with the GK model. When we applied the GB model in DAP-LW, the PA was 0.82 for MDs, 0.83 for MDe and 0.84 for MM, and in AP-LW, it ranged from 0.80 for MDs to 0.81 for MDe and MM (Figure 3). For the WW environment in AP the model-method combinations presented the same values, with a PA of 0.83 for both the GK and GB (Figure 3). For WW the PA ranged from 0.86 for MM to 0.87 for MDe and MDs, respectively, for both the GK and GB (Figure 3).

### 3.6 EGG

For rubber tree breeding, the investigated alternative breeding strategies differed considerably in the number of years required to finish one breeding cycle. For classic improvement, we consider a minimum time of ten years for the beginning of the selection of the best genotypes. In the case of GS, we consider three years for the first selection.

To determine the best EGG, we used the best scenario found in the analyses, so we chose to work with the GK matrix and the phenotypic data from the WW environment; however, we present the other results in Supplementary Table 2. Using the classical breeding method (CBM), which takes into account only the phenotypic data, the selection gains ranged from 0.058 (DAP) to 0.064 (AP) (Figure 4). When we predict breeding value without environmental effects (BSG), genetic gains increase from 0.068 in AP to 0.114 in DAP (figure 4). When we incorporated the molecular information in a single environment (SM), the genetic gain increased to 0.160 (DAP) and 0.105 (AP). When we used the multi-environment strategy, the gains were much higher; for MM, the genetic gain was 0.497 for DAP and 0.434 for AP; in MDs and MDe, we obtained 0.503 and 0.434 for both DAP and AP, respectively (Figure 4).

**Figure 4.**
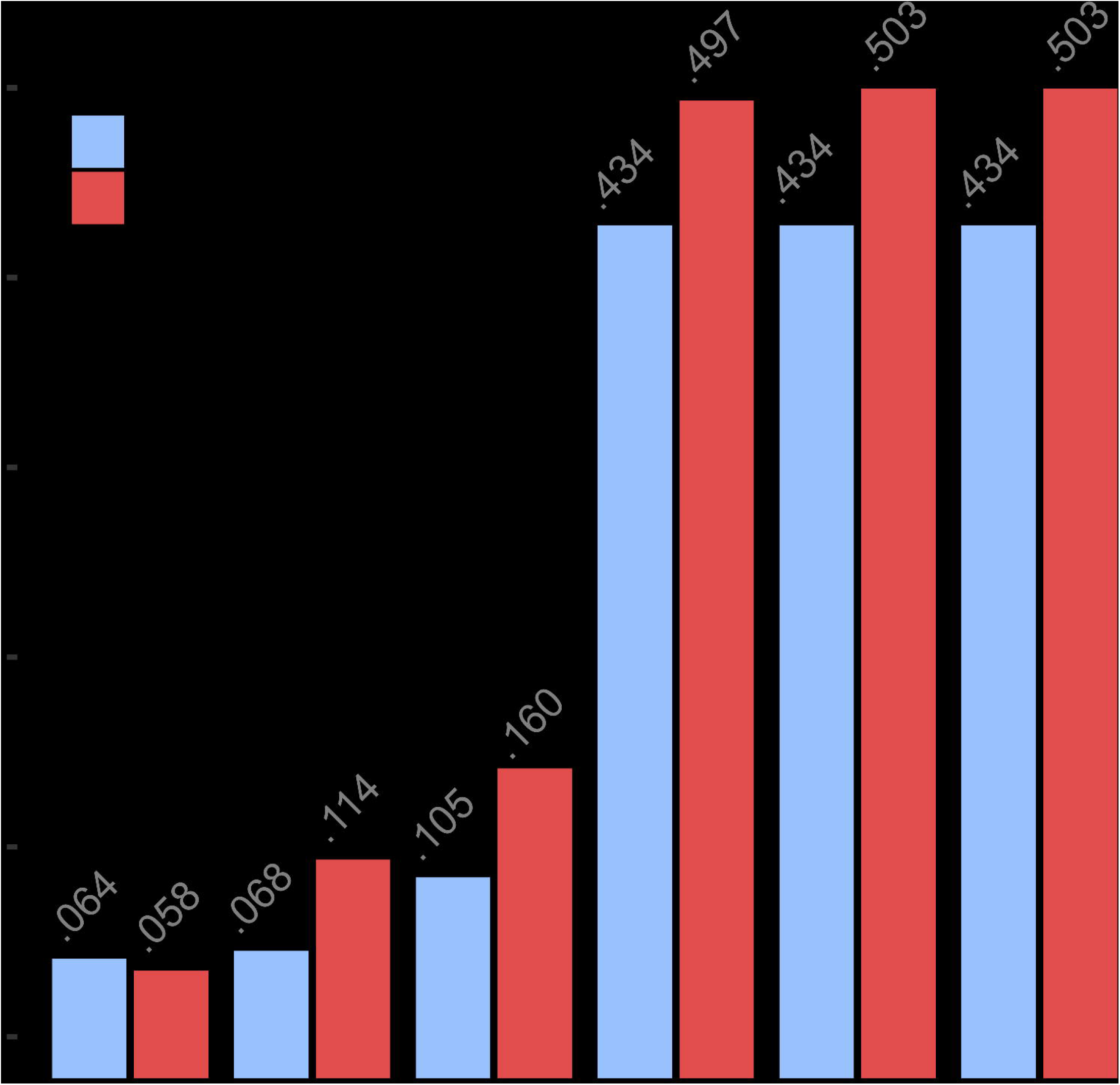
Expected genetic gain (EGG) using the classical breeding method (CBM) and genomic prediction without multi-environmental data (BSG), with the single-environment, main genotypic effect model (SM), with the multi-environment, genotypic effect model (MM), with the multi-environment, single variance G×E model (MDs), and with the multi-environment, environment-specific variance G×E model (MDe) with the GK kernel method in the WW environment.

## 4 Discussion

In recent years, many statistical models have been proposed for applying GS in plant and animal breeding programs. However, to the best of our knowledge, these models have not yet been applied to rubber tree breeding programs.

The efficiency of early selection depends mainly on the early-mature correlation and the heritability of juvenile traits. In this study, all pairwise correlations between environments were strong and positive. This finding is important because the G×E model has the limitation of better and more efficient prediction when applied to subsets of environments that have positive and similar correlations (Crossa et al., 2016).

In a study by Moreti et al. (1994), with estimates of genetic parameters and expected gains with the selection of juvenile characters in rubber tree progenies using classical breeding, some parameters stood out positively (rubber production, bark thickness, and stem circumference). Gonçalves et al. (1996) observed the same behavior of the results obtained by Moreti et al. (1994), showing a correlation and its applicability in the selection process. Strong phenotypic and genetic correlations were observed between yield and stem diameter, indicating the possibility of obtaining young clones of good productive capacity and great vigor (Gonçalves et al., 1984). Plants with rapid development of trunk circumference may be more productive, and this may be a useful feature with which to predict more productive hybrids via GS, together with the fact that latex production has a good heritability, better than growth in diameter, because the influence of the rootstock is lower in the production, this will be very important in future studies with this population.

The multi-environment genomic prediction was successfully implemented using a GBLUP model; however, depending on the genetic architecture of the trait and germplasm, nonlinear semiparametric approaches, such the GK, could produce better accuracies (Cuevas et al., 2016). We compared two models that used the GB and GK methods, and similar values were found for the PA in all tested conditions.

Using genomic prediction without multi-environment data (BSG) for both GB and GK produced the same results (0.18) for AP and a small change from 0.26 to 0.27 between GK and GB, respectively. When a single-environment model was applied to only one environment (LW or WW), the best PA values were in the environment of higher water availability. For AP, we obtained the best values (0.20) using the GK matrix. On the other hand, for DAP, the best PA was obtained via GK matrix (0.28) (Figure 2).

In this study, the PA of multi-environment models was assessed by applying the CV strategy. The CV2 validation strategy performed better than the CV1 strategy when applied to the multi-environment models (Lopez-Cruz et al., 2015; Crossa et al., 2016); this result was expected because we did not evaluate all individuals in both environments. Considering only random CV2, the PA was slightly higher in WW conditions for both characters (AP and DAP), ranging from 0.87 (MDs-GK-WW) to 0.81 (GK-MDe-LW).

Evaluating plants during the season with the best hydric conditions resulted in a higher PA, which may have occurred because more measurements were taken in WW than in LW, which in turn improved the performance of the analyses. However, according to (Conson et al., 2018), the climatological water balance revealed that phenotypic sampling was performed in consecutive water deficit periods, with exceptions including very brief intervals with very high precipitation levels, and the two regions where the experiments were carried out suffer from low water availability for a large part of the year.

Multi-environment models are superior to single-environment genomic models with the GBLUP and GK. This finding suggests that introducing interactions between markers and environmental conditions can increase the proportion of variance accounted for by the model and, more importantly, can increase the PA. GxE interactions are essential in many aspects of a breeding program, and the increase in PA with the inclusion of environmental information represents a favorable result with important implications for both breeding and agronomic recommendations. Interactions in field trials affect mature selection and early selection; thus, when evaluating the effectiveness of early selection, it is imperative to determine whether the GxE interaction among sites has a meaningful impact on the early-mature genetic correlation.

The application of eight combinations of four models (SM, MM, MDs, and MDe) and two kernel methods (GBLUP and GK) to rubber tree data sets showed that models with the nonlinear GK had slightly higher PAs than the models with the linear GBLUP kernel. According to Gianola et al. (2014), the GK has a better predictive ability and a more flexible structure than the GBLUP, and the GK can capture nonadditive effects between markers.

Akdemir and Jannink (2015) presented different choices for estimating kernel functions: linear kernel matrices incorporate only the additive effects of the markers, polynomial kernels incorporate different degrees of marker interactions, and the GK function uses complex epistatic marker interactions. GKs would be more appropriate for GS for rubber trees because of the possibility of exploiting these local epistatic effects captured in the GK and their interaction with environments.

Many GS studies in plants have focused on breeding programs that generally evaluate crops in multiple environments, such as in different seasons/years or geographic locations, to determine performance stability across environments (Crossa et al., 2016) and identify markers whose effects are stable across environments as well as those that are environment-specific (Crossa et al., 2016; Oakey et al., 2016). Lopez-Cruz et al. (2015) extended the single-trait GBLUP model to a multi-environment context and showed substantial gains in PA with the multi-environment model relative to single-environment analysis in wheat.

The advantages of GS applied to the improvement of forest species have been successfully demonstrated. For example, Wong and Bernardo (2008) and Iwata et al. (2011) demonstrated the potential uses of GS, and all concluded that it could radically increase tree breeding efficiency. The advantage of marker-based relationship matrices is that gaps in pairwise relatedness in forest tree pedigrees are filled, which leads to an increase in the accuracy of selecting breeding candidates (Muller et al., 2017; Tan et al., 2017).

Cuevas et al. (2016) modeled G×E interactions using both genetic markers and environmental covariates, and (Granato et al., 2018) introduced the Bayesian Genomic Genotype × Environment (BGGE) R package, which fits genomic linear mixed models to single environments and multi-environments with GE models. These studies showed that modeling multiple-environment interactions can lead to substantial gains in the PA of GS for rubber tree breeding programs.

GS is expected to increase the accuracy of selection, especially for traits that cannot be measured directly from breeding candidates and for traits with low heritabilities (Meuwissen et al., 2001), which was confirmed in this study: the selection gain with GS varied between 0.434 and 0.503, while the genetic gain with classical breeding varied between 0.058 and 0.064 for AP and DAP, respectively. Comparing the conventional model of genetic breeding (CBM) for rubber trees with the use of GS applying the multi-environment strategy (MM, MDe and MDS), GS produced a genetic gain 6.7 and 8.7 times higher for AP and DAP, respectively. GS also resulted in a more balanced selection response in the two traits (DAP and AP) and thus is preferred over traditional selection because of the time saved in the selection of superior genotypes.

Developing new rubber tree cultivars adaptable to non-traditional rubber growing regions is fundamental for the success of rubber tree plantations. Cultivars considered productive in some regions of Brazil may behave differently in other areas of the same region, especially areas with different edaphoclimatic characteristics. Several agroclimatic elements, such as prolonged low temperature and low precipitation in the winter, are the major factors limiting the development and production of the rubber tree and contribute to a large amount of variability in the behavior of cultivars (Ortolani et al., 1996).

In rubber trees, the time required to complete a breeding cycle and recommend a clone for commercial production can span multiple decades and is mainly divided into three selection stages. First, the aim is to obtain progenies by controlled or open pollination to establish nurseries. At two and a half years, based on yield evaluations performed by early testing of yield, vigor, and tolerance to diseases, the breeding plants are selected and cloned for testing at a small scale. In this second stage of the selection cycle, after the first two years of tapping, promising clones are multiplied and subsequently evaluated in large-scale or regional trials. This last stage usually takes some 12 to 15 years, until it is possible to recommend a clone for large-scale cropping. It, therefore, takes approximately 30 years to complete the breeding cycle, from controlled pollination to final cultivar recommendation (Gonçalves and Fontes, 2012). The use of GS could dramatically reduce the time required for completion of a cycle of genetic improvement by eliminating progeny phenotypic testing aimed at selecting the best individuals (replaced by GS), significantly accelerating the genetic gain relative to that obtained by classical breeding. Another advantage of GS is that more candidate genotypes are generated; therefore, the population size for selection as well. All of these candidates are genotyped, and those with the best-predicted test cross values are evaluated in the field, which can be regarded as an indirect selection.

With the rapid advances in and declining costs of genotyping methods, balanced against the overall costs of managing large progeny trials and the potential for increased gains per unit time, our cautiously optimistic expectation is that GS has excellent potential to be implemented in rubber tree breeding programs. However, further studies examining populations with a different structure (which were not assessed in this initial work) are necessary before recommending GS for operational implementation in tree breeding programs.

## Author contributions

LS, FF, PG, RN and AS designed the study and performed the experiments; LS, FF, and RN analyzed the data; and LS, FF and AS wrote the manuscript.

## References

Akdemir, D., and Jannink, J.L. (2015). Locally epistatic genomic relationship matrices for genomic association and prediction. Genetics 199, 857–871.

Bandeira, E.S.M., Cuevas, J., De Oliveira Couto, E.G., Perez-Rodriguez, P., Jarquin, D., Fritsche-Neto, R., et al. (2017). Genomic-enabled Prediction in maize using kernel models with genotype × environment interaction. G3 (Bethesda) 7, 1995–2014.

Bartholome, J., Van Heerwaarden, J., Isik, F., Boury, C., Vidal, M., Plomion, C., et al. (2016). Performance of genomic prediction within and across generations in maritime pine. BMC Genomics 17, 604.

Burgueño, J., De Los Campos, G., Weigel, K., and Crossa, J. (2012). Genomic prediction of breeding values when modeling genotype × environment interaction using pedigree and dense molecular markers. Crop Sci. 52, 707–719.

Conson, A. R. O., Taniguti, C. H., Amadeu, R. R., Andreotti, I. A. A., de Souza, L. M., Dos Santos, L. H. B., et al. (2018). High-resolution genetic map and QTL analysis of growth-related traits of Hevea brasiliensis cultivated under suboptimal temperature and humidity conditions. Front. Plant Sci. 9:1255. doi: 10.3389/fpls.2018.00513.

Crossa, J., De Los Campos, G., Maccaferri, M., Tuberosa, R., Burgueño, J., and Pérez-Rodríguez, P. (2016). Extending the marker × environment interaction model for genomic-enabled prediction and genome-wide association analysis in durum wheat. Crop Sci. 56, 2193–2209.

Cuevas J, Crossa J, Soberanis V, Pérez-Elizalde S, Pérez-Rodríguez P, de los Campos G, et al. (2016). Genomic prediction of genotype × environment interaction kernel regression models. Plant Genome. 9:3.

Danecek, P., Auton, A., Abecasis, G., Albers, C.A., Banks, E., Depristo, M.A., et al. (2011). The variant call format and VCFtools. Bioinformatics 27, 2156–2158.

El-Dien O.G., Ratcliffe B, Klápště J, Chen C, Porth I, El-Kassaby YA. (2015). Prediction accuracies for growth and wood attributes of interior spruce in space using genotyping-by-sequencing. BMC Genomics. 16:370.

Furlani, R.C.M., Moraes, M.L.T.D., Resende, M.D.V.D., Furlani Junior, E., Gonçalves, P.D.S., Valério, W.V.F, et al. (2005). Estimation of variance components and prediction of breeding values in rubber tree breeding using the REML/BLUP procedure. Genet. Mol. Biol. 28, 271–276.

Gianola, D., Weigel, K.A., Krämer, N., Stella, A., and Schön, C.C. (2014). Enhancing genome-enabled prediction by bagging genomic BLUP. PLoS One 9, e91693.

Glaubitz, J.C., Casstevens, T.M., Lu, F., Harriman, J., Elshire, R.J., Sun, Q., et al. (2014). TASSEL-GBS: a high capacity genotyping by sequencing analysis pipeline. PLoS One 9, e90346.

Gonçalves, P.S., and Fontes, J.R.A. (2012). “Domestication and breeding of rubber tree,” in Domestication and Breeding – Amazonian Species, eds. A. Borém, M.T.G. Lopes, C.R.C. Clement & H. Noda (Viçosa: Suprema Editora Ltda), 393–419.

Gonçalves, P.S., Martins, A.L.M., Bortoletto, N., and Tanzini, M.R. (1996). Estimates of genetic parameters and correlations of juvenile characters basead on open pollinated progenies of Hevea. Rev. Bras. Genet. 19, 105–111.

Gonçalves, P.S., Rossetti, A.G., Valois, A.C.C., and Viegas, I.J. (1984). Genetic and phenotypic correlations between some quantitative traits in juvenile clonal rubber trees (Hevea spp.). Rev. Bras. Genet. II, 95–107.

Gonçalves, P.S., Silva, M.A., Gouvêa, L.R.L., and Scaloppi-Junior, E.J. (2006). Genetic variability for girth growth and rubber yield traits in *Hevea brasiliensis*. Sci. Agric. 63, 246–254.

Granato, I., Cuevas, J., Luna-Vazquez, F., Crossa, J., Montesinos-Lopez, O., Burgueno, J., et al. (2018). BGGE: a new package for genomic-enabled prediction incorporating genotype × environment interaction models. G3 (Bethesda) 8, 3039–3047.

Granato, I., and Fritsche-Neto, R. (2018). snpReady: Preparing genotypic datasets in order to run genomic; analysis. R package version 0.9.6. Available: https://CRAN.R-project.org/package=snpReady.

Habier, D., Fernando, R.L., and Dekkers, J.C. (2007). The impact of genetic relationship information on genome-assisted breeding values. Genetics 177, 2389–2397.

Isik, F. (2014). Genomic selection in forest tree breeding: the concept and an outlook to the future. New For. 45, 379–401.

Isik, F., Bartholome, J., Farjat, A., Chancerel, E., Raffin, A., Sanchez, L., et al. (2016). Genomic selection in maritime pine. Plant Sci. 242, 108–119.

Iwata, H., Hayashi, T., and Tsumura, Y. (2011). Prospects for genomic selection in conifer breeding: a simulation study of *Cryptomeria japonica*. Tree Genet. Genomes 7, 747–758.

Jarquin, D., Crossa, J., Lacaze, X., Du Cheyron, P., Daucourt, J., Lorgeou, J., et al. (2014). A reaction norm model for genomic selection using high-dimensional genomic and environmental data. Theor. Appl. Genet. 127, 595–607.

Krchov, L.-M., and Bernardo, R. (2015). Relative efficiency of genomewide selection for testcross performance of doubled haploid lines in a maize breeding program. Crop Sci. 55, 2091–2099.

Langmead, B., and Salzberg, S.L. (2012). Fast gapped-read alignment with Bowtie 2. Nat. Methods 9, 357–359.

Lima, B.M. (2014). Bridging genomics and quantitative genetics of Eucalyptus: genome-wide prediction and genetic parameter estimation for growth and wood properties using high-density SNP data. [thesis]. [Piracicaba, Brazil]: University of São Paulo.

Lopez-Cruz, M., Crossa, J., Bonnett, D., Dreisigacker, S., Poland, J., Jannink, J.L., et al. (2015). Increased prediction accuracy in wheat breeding trials using a marker × environment interaction genomic selection model. G3 (Bethesda) 5, 569–582.

Lorenz, A.J., Chao, S., Asoro, F.G., Heffner, E.L., Hayashi, T., Iwata, H., et al. (2011). “Chapter Two – Genomic selection in plant breeding: knowledge and prospects,” in Advances in Agronomy, ed. D.L. Sparks. (San Diego: Academic Press), 77–123.

Meuwissen, T.H., Hayes, B.J., and Goddard, M.E. (2001). Prediction of total genetic value using genome-wide dense marker maps. Genetics 157, 1819–1829.

Muller, B.S.F., Neves, L.G., De Almeida Filho, J.E., Resende, M.F.R., Jr., Munoz, P.R., Dos Santos, P.E.T., et al. (2017). Genomic prediction in contrast to a genome-wide association study in explaining heritable variation of complex growth traits in breeding populations of *Eucalyptus*. BMC Genomics 18, 524.

Munõz, F., and Sanchez, L. (2017). breedR: statistical methods for forest genetic; resources analysts. R package version 0.12-2. Available: https://github.com/famuvie/breedR.

Oakey, H., Cullis, B., Thompson, R., Comadran, J., Halpin, C., and Waugh, R. (2016). Genomic selection in multi-environment crop trials. G3 (Bethesda) 6, 1313–1326. doi: 10.1534/g3.116.027524/-/DC1

Ortolani, A., Sentelhas, P., Camargo, M., Pezzopane, J., and Gonçalves, P. (1996). Agrometeorological models to estimate annual and seasonal production of latex in rubber. Revista Brasileira de Agrometeorologia 4, 147–150.

Pérez-Elizalde, S., J. Cuevas, P. Pérez-Rodríguez, and J. Crossa (2015). Selection of the Bandwidth Parameter in a Bayesian Kernel Regression Model for Genomic-Enabled Prediction. J. Agric. Biol. Environ. Stat. 20(4): 512–532.

Pérez-Rodríguez, P., Crossa, J., Bondalapati, K., De Meyer, G., Pita, F., and Campos, G.D.L. (2015). A pedigree-based reaction norm model for prediction of cotton yield in multienvironment trials. Crop Sci. 55, 1143–1151.

Ratcliffe, B., El-Dien, O.G., Klápště, J., Porth, I., Chen, C., Jaquish, B., et al. (2015). A comparison of genomic selection models across time in interior spruce (Picea engelmannii × glauca) using unordered SNP imputation methods. Heredity (Edinb) 115, 547–555.

Resende, M.D., Resende, M.F., Jr., Sansaloni, C.P., Petroli, C.D., Missiaggia, A.A., Aguiar, A.M., et al. (2012a). Genomic selection for growth and wood quality in *Eucalyptus*: capturing the missing heritability and accelerating breeding for complex traits in forest trees. New. Phytol. 194, 116–128.

Resende, M.F., Jr., Munoz, P., Acosta, J.J., Peter, G.F., Davis, J.M., Grattapaglia, D., et al. (2012b). Accelerating the domestication of trees using genomic selection: accuracy of prediction models across ages and environments. New. Phytol. 193, 617–624.

Souza, L.M., Gazaffi, R., Mantello, C.C., Silva, C.C., Garcia, D., Le Guen, V., et al. (2013). QTL mapping of growth-related traits in a full-sib family of rubber tree (Hevea brasiliensis) evaluated in a sub-tropical climate. PLoS One 8, e61238.

Tan, B., Grattapaglia, D., Martins, G.S., Ferreira, K.Z., Sundberg, B., and Ingvarsson, P.K. (2017). Evaluating the accuracy of genomic prediction of growth and wood traits in two *Eucalyptus* species and their F1 hybrids. BMC Plant Biol. 17, 110.

Tang, C., Yang, M., Fang, Y., Luo, Y., Gao, S., Xiao, X., et al. (2016). The rubber tree genome reveals new insights into rubber production and species adaptation. Nat. Plants 2, 16073.

Wong, C.K., and Bernardo, R. (2008). Genomewide selection in oil palm: increasing selection gain per unit time and cost with small populations. Theor. Appl. Genet. 116, 815–824.

Zapata-Valenzuela, J., Whetten, R.W., Neale, D., Mckeand, S., and Isik, F. (2013). Genomic estimated breeding values using genomic relationship matrices in a cloned population of loblolly pine. G3 (Bethesda) 3, 909–916.

