## Supplementary material for "Genomic Selection in Rubber Tree Breeding: A Comparison of Models and Methods for dealing with G × E"

**Supplementary Tables**

**Supplementary Table 1.** Dates of the phenotyping of the two populations.

|  | Michelin | | IAC | |
| --- | --- | --- | --- | --- |
| year | LW | WW | LW | WW |
| 1 | October/07 | - | December/13 | June/13 |
| 2 | October/08 | April/08 | November/14 | May/14 |
| 3 | October/09 | April/09 | - | June/15 |
| 4 | October/10 | April/10 | - | June/16 |

**Supplementary Table 2.** Expected genetic gain (EGG) using the classical breeding method (CBM) or genomic prediction without multienvironmental data (BSG), with the single-environment, main genotypic effect model (SM), with the multienvironment, genotypic effect model (MM), with the multienvironment, single variance G×E model (MDs), and with the multienvironment, environment-specific variance G×E model (MDe) with the GBLUP kernel method in the WW environment.

| Models | Character | Environment | Matrix | EGG |
| --- | --- | --- | --- | --- |
| CBM | DAP | - | - | 0.077 |
|  | AP | - | - | 0.087 |
|  | DAP | LW | - | 0.058 |
|  | DAP | WW | - | 0.058 |
|  | AP | LW | - | 0.065 |
|  | AP | WW | - | 0.064 |
| BSG | DAP | - | GK | 0.114 |
|  | DAP | - | GB | 0.115 |
|  | AP | - | GK | 0.068 |
|  | AP | - | GB | 0.074 |
| SM | AP | WW | GK | 0.105 |
|  | AP | WW | GB | 0.100 |
|  | AP | LW | GK | 0.092 |
|  | AP | LW | GB | 0.087 |
|  | DAP | WW | GK | 0.098 |
|  | DAP | WW | GB | 0.162 |
|  | DAP | LW | GK | 0.109 |
|  | DAP | LW | GB | 0.109 |
| MM | DAP | LW | GK | 0.480 |
|  | DAP | LW | GB | 0.480 |
|  | DAP | WW | GK | 0.497 |
|  | DAP | WW | GB | 0.497 |
|  | AP | LW | GK | 0.420 |
|  | AP | LW | GB | 0.415 |
|  | AP | WW | GK | 0.434 |
|  | AP | WW | GB | 0.434 |
| MDs | DAP | LW | GK | 0.480 |
|  | DAP | LW | GB | 0.475 |
|  | DAP | WW | GK | 0.503 |
|  | DAP | WW | GB | 0.503 |
|  | AP | LW | GK | 0.415 |
|  | AP | LW | GB | 0.415 |
|  | AP | WW | GK | 0.434 |
|  | AP | WW | GB | 0.434 |
| MDe | DAP | LW | GK | 0.480 |
|  | DAP | LW | GB | 0.469 |
|  | DAP | WW | GK | 0.503 |
|  | DAP | WW | GB | 0.503 |
|  | AP | LW | GK | 0.415 |
|  | AP | LW | GB | 0.410 |
|  | AP | WW | GK | 0.434 |
|  | AP | WW | GB | 0.434 |

**Supplementary Figures**

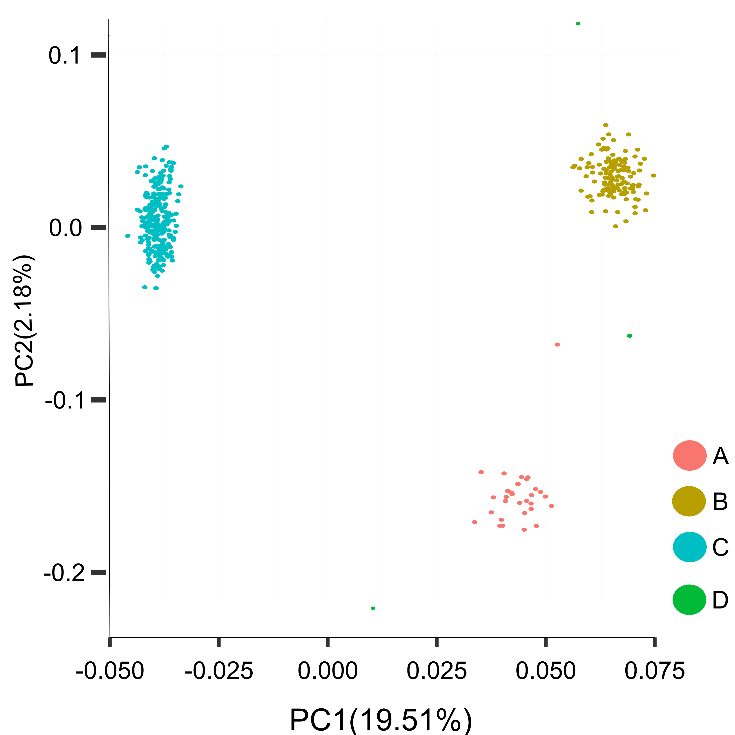

**Supplementary Figure 1** Population structure analysis of 411 rubber tree genotypes with the first two principal components (PCs).

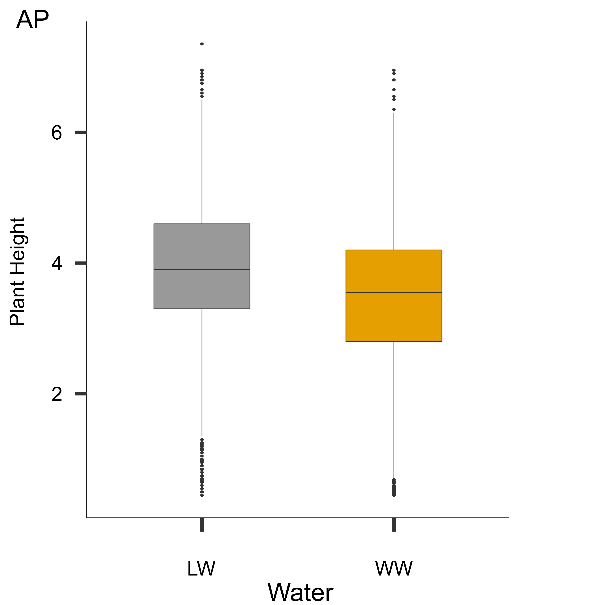

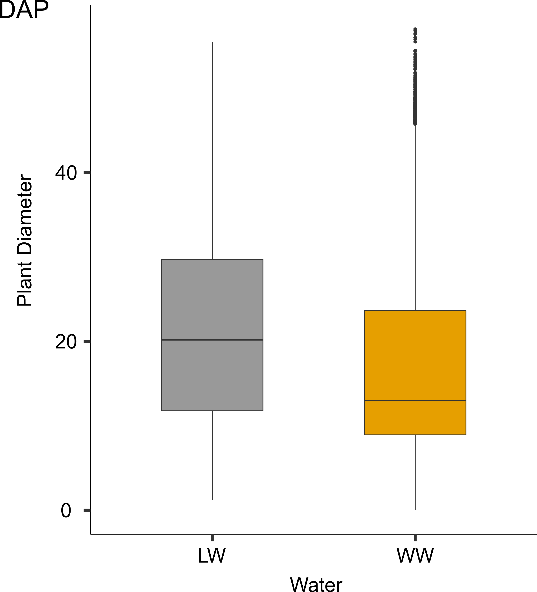

**Supplementary Figure 2**. Box plot of plant height (PA) and plant diameter (DAP) in the two environments: WW (well-watered) and LW (low water).

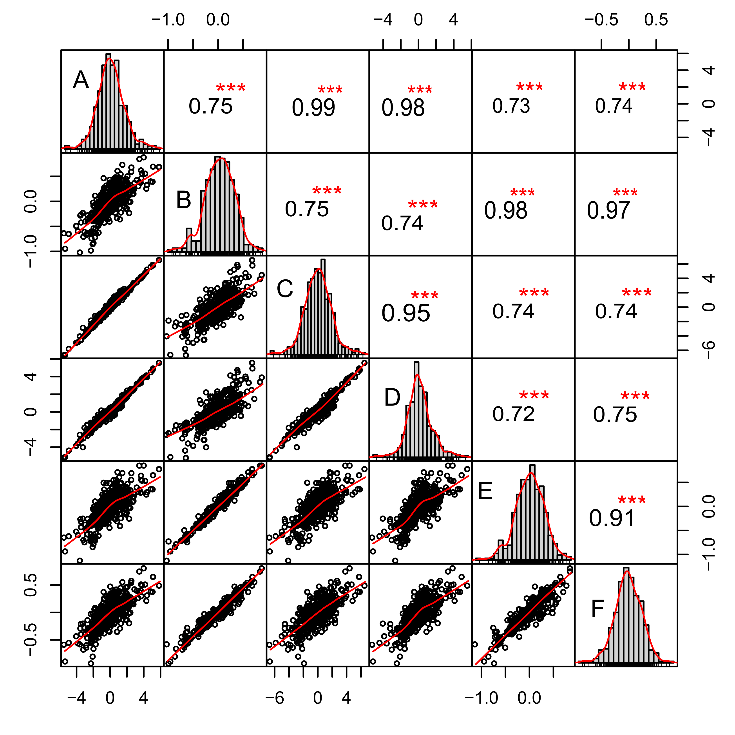

**Supplementary Figure 2.** Estimates of genetic correlations between AP (plant height) and DAP (stem diameter) in LW (low-water) and WW (well-watered) conditions.

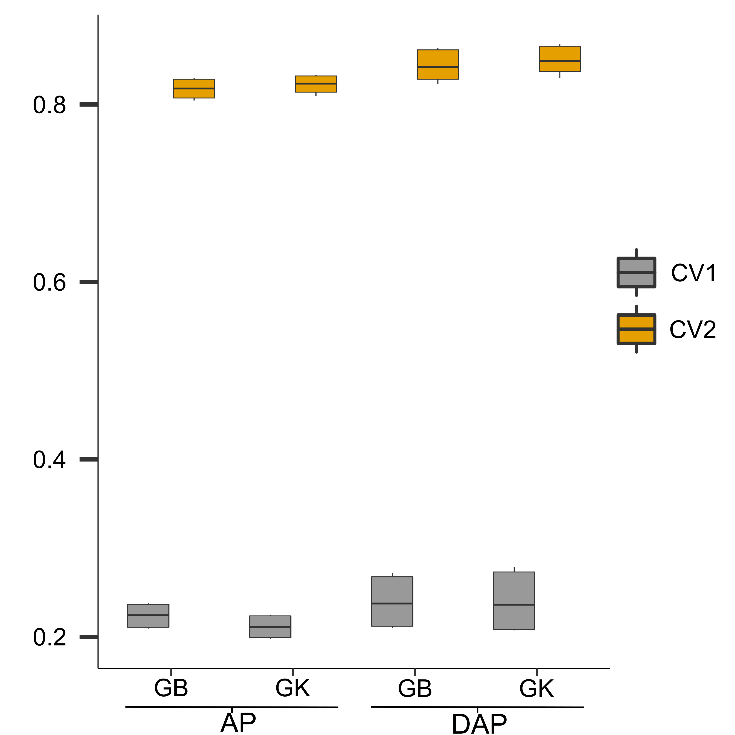

**Supplementary Figure 3.** Mean prediction accuracies (PAs) for the diameter (DAP) and plant height (PA) of rubber trees using the genomic best linear unbiased prediction (GBLUP) and Gaussian kernel (GK) methods under the cross-validation (CV) scenarios CV1 and CV2.
